# Relative Brain Age Is Associated with Socioeconomic Status and Anxiety/Depression Problems in Youth

**DOI:** 10.1101/2022.09.15.505331

**Authors:** Jacob W. Cohen, Bruce Ramphal, Mariah DeSerisy, Yihong Zhao, David Pagliaccio, Stan Colcombe, Michael P. Milham, Amy E. Margolis

**Author notes:** **Author Note**. This study was not pre-registered.

## Abstract

Socioeconomic status (SES) has been linked to differences in brain structure and psychiatric risk across the lifespan. Despite many neuropsychiatric disorders emerging in childhood, few studies have examined the influence of SES on brain aging and psychopathology in youth.

We re-analyzed relative brain age (RBA) data from the Healthy Brain Network to examine the influence of SES components (parent education, occupation, household income-to-needs ratio (INR), public assistance enrollment) on RBA. RBA was previously determined using covariation patterns for cortical morphology, white, and subcortical gray matter volumes without SES in predictive models. We also examined associations between RBA and psychiatric symptoms (child behavior checklist). Full case analysis included 470 youth (5-17 years; 61.3% male), self-identifying as White (55%), African American (15%), Hispanic (9%), or multiracial (17.2%). Mean household income was 3.95±2.33 (Mean±SD) times the federal poverty threshold. Multiple linear regression examined if 1) SES components associated with RBA, and 2) RBA associated with psychiatric symptoms. Models covaried for sex, scan location, and parent psychiatric diagnoses.

RBA associated with public assistance (p = 0.03), parent occupation (p = 0.01), and parent psychiatric diagnosis (p = 0.01), but not with INR and parent education. Parent occupation (p = 0.02) and RBA (p = 0.04) associated with CBCL anxiety/depression scores.

Components of SES associated with brain aging, underscoring the risk of omitting these factors in developmental brain research. Further, delayed brain aging was associated with low parental occupational prestige and child anxiety/depression scores, suggesting a possible biological pathway from SES to mental health risk.

## INTRODUCTION

Living in lower SES contexts confers an increased risk for receiving a psychiatric diagnosis (Merikangas et al., 2010) and predicts higher rates of internalizing problems (Amone-P’Olak et al., 2011) in adolescence. These associations endure beyond adolescence; lower childhood SES predicts more anxiety symptoms (Lähdepuro et al., 2019) and anxiety disorder diagnoses (McLaughlin et al., 2010) in adulthood. Further, lower childhood SES associates with lifetime depression risk (Gilman et al., 2003; Power et al., 2007), episode onset (Lorant et al., 2003), and persistence (Lorant et al., 2003; Power et al., 2007). SES is often measured as a combination of ostensibly interchangeable components (e.g., parent education, occupation, income, public assistance). Examining these components independently, however, shows that childhood financial hardship robustly predicts mental health later in life while parent education, occupational class and income do not (Lahelma et al., 2006). Similarly, public assistance predicts onset of all classes of mental health disorders while parent education predicts persistence and severity of the condition, with variable effect sizes across different classes of disorders (McLaughlin et al., 2011). Specifically, socioeconomic differences in depression prevalence, incidence and persistence vary with how SES is measured (Lorant et al., 2003). Though SES components may be strongly correlated, differences in their associations across metrics of mental health suggest they capture distinct aspects of environmental and genetic variation, and warrant individual attention (Braveman et al., 2005; Duncan & Magnuson, 2012).

Childhood SES has also been linked to differences in brain structure. Specifically, lower household SES associates with lower total gray matter volume (Dennis et al., 2022; Jednoróg et al., 2012; Luby et al., 2013; Mackey et al., 2015; McDermott et al., 2019), cortical gray matter volume (Luby et al., 2013; Mackey et al., 2015), cortical thickness (Colich et al., 2020; Mackey et al., 2015; McDermott et al., 2019; Noble et al., 2015), and cortical surface area (McDermott et al., 2019) in youth. Regionally, SES is positively associated with hippocampal (Botdorf et al., 2022; Hanson et al., 2011; Jednoróg et al., 2012; Luby et al., 2013; McDermott et al., 2019; Noble et al., 2015, 2012; Ramphal et al., 2021) and amygdala (Assari, 2020; Assari et al., 2020; Luby et al., 2013; McDermott et al., 2019; Ramphal et al., 2021) volumes across childhood and adolescence, although some studies have failed to replicate findings in the amygdala (Hanson et al., 2011; Noble et al., 2012). Lower SES has also been shown to associate with lower global white matter volumes in one study (McDermott et al., 2019) but not in others (Jednoróg et al., 2012; Mackey et al., 2015). In line with differing associations between SES components and mental health outcomes, parent education and income differ in their ability to predict cortical thickness across the brain (Lawson et al., 2013; Noble et al., 2015, 2012; Rakesh et al., 2022) as well as hippocampal volume (Noble et al., 2015). These findings suggest different SES components may be biologically distinct and could help explain how SES components result in divergent mental health outcomes.

The association between SES and the structure of the child and adolescent brain has been widely examined, but synthesizing these findings into clinically useful information is difficult, given the number of regions involved, heterogeneous patterns of development across the brain, and heterogeneity across neuroimaging modalities (e.g., fMRI, diffusion tensor imaging; Erus et al., 2015). Machine learning methods offer new opportunities to synthesize complex, high-dimensional imaging data and characterize normative developmental trajectories. In one application, machine learning is leveraged to predict an individual’s age based on many structural brain measures. Differences between an individual’s model predicted age and their actual chronological age, termed brain age gap, have been examined, such that positive gaps represent accelerated aging and negative gaps represent delayed aging (Franke & Gaser, 2019). Such measures may be markers of psychopathology; accelerated aging associates with post-traumatic stress disorder (Clausen et al., 2022), schizophrenia (Hajek et al., 2019; Lee et al., 2021; Nenadić et al., 2017; Shahab et al., 2019; J. Wang et al., 2021), depression (Ballester et al., 2021; Dunlop et al., 2021; Han et al., 2021; Tang et al., 2021) and psychosis (Kolenic et al., 2018; McWhinney et al., 2021) in adults. Brain age studies in youth are relatively limited and it is not yet clear which direction (i.e., accelerated or delayed) might portend risk. In youth, accelerated aging has been shown to associate with greater general psychopathology (Cropley et al., 2021) and, specifically, psychosis (Chung et al., 2018), obsessive compulsive disorder (Cropley et al., 2021), and depression (Drobinin et al., 2021). Delayed aging has been shown to associate with greater autism symptom burden (Q. Wang et al., 2021) and contrary to the aforementioned studies, general psychopathology (Luna et al., 2021; Lund et al., 2022). Less is known, however, about factors that contribute to the brain age gap. Although household SES has not been specifically examined in relation to brain age, accelerated brain aging has been shown to associate with greater childhood physical neglect (Keding et al., 2021), emotional/sexual abuse and exposure to violence (Drobinin et al., 2021), as well as neighborhood disadvantage in early adolescence with inverse findings in late adolescence (Rakesh, Cropley, et al., 2021).

In the present study, we reanalyze prior validated brain age measurements in children in the Healthy Brain Network (HBN) sample with an interest in understanding the association between household SES and the brain age gap (based on structural brain morphometry) across childhood and adolescence (age 5-17 years). Because this work is intended as a follow-up to the prior study, subjects were limited to those included in the prior study. We parse SES into specific components including income-to-needs ratio (INR), parent education, parent occupation, and public assistance utilization in the same model to account for shared variance. We hypothesized that these specific components of SES would be distinctly associated with brain aging; the expected direction of this effect remains unclear due to equivocal findings in the literature (Drobinin et al., 2021; Keding et al., 2021; Rakesh, Cropley, et al., 2021). Second, we explore links between brain age and concurrent parent-related problem behaviors to ground our findings in the literature linking brain age to mental health outcomes.

## METHODS

### Participants

The current study was intended to further examine effects of SES components on outcomes from an initial study of brain aging in youth in HBN (Zhao et al., 2019), an ongoing research initiative aimed at collecting freely available imaging data from a community sample of New York City youth enriched for psychiatric problems (Alexander et al., 2017). The initial study leveraged data collected between the years of 2015 and 2018. Subjects were excluded when there was an immediate safety concern, a cognitive or behavioral impairment that impeded data collection, or medical/psychiatric conditions that would confound future research with the data (Alexander et al., 2017). Briefly, participants underwent MRI imaging and environmental, behavioral and psychological phenotypic assessment. MRI scans were conducted at 3 sites: Rutgers University Brain Imaging Center, Citigroup Biomedical Imaging Center and a mobile scanner placed in Staten Island. Our analyses included 470 of the original 869 participants training set, who also had available and complete demographic, socioeconomic, and phenotypic data for a full case analysis. Written consent was obtained from a legal guardian and written assent obtained from all participants.

### Relative Brain Age (RBA)

In prior work (Zhao et al., 2019), MRI data was preprocessed using the FreeSurfer software (Dale et al., 1999), shape measures were extracted using the Mindboggle software package (Klein et al., 2017), and machine learning was leveraged to predict an individual’s age based on structural brain morphometry. The training sample consisted of N = 869 (530 male, 339 female, mean age = 10.54, age range: 5.02 – 17.95 years) structural brain images from the HBN biobank. Cortical mean curvature, white matter volume, cortical thickness, surface area, gray matter volume, and travel depth were measured for 62 regions of interest (Desikan et al., 2006) and gray matter volume estimates were also measured for an additional 16 subcortical ROIs. To best capture the covariation across shape measures and distinct to each measure across ROIs, a data reduction technique called Joint and Individual Variance Explained (JIVE) was implemented (Lock et al., 2013). Joint and individual components, representing covariation patterns shared across and/or specific to individual shape measure and total volumes, as well as the total volumes of the intracranium, brain stem, cerebrospinal fluid, and subcortical gray matter, were entered into a ridge regression to build a highly predictive model of chronological age (Mean Absolute Error =1.41 years, adjusted R_2_=.71; Zhao et al., 2019).

In regression modeling there is a prediction bias to the mean, that manifests in brain age research as an overestimation of predicted age in younger participants, and an underestimation in older participants (Smith et al., 2019). In the current study, instead of using the difference in predicted and actual age to measure aging, relative brain age (RBA) was computed by regressing predicted age on chronological age in N = 869 participants and saving the residuals in order to eliminate this bias (Ning et al., 2020; Rokicki et al., 2021) and henceforth providing us with an implicit control for age in all subsequent analyses. Positive RBA values are interpreted as accelerated brain aging and negative RBA values, as delayed aging.

### Socioeconomic Status Components

Data on parent educational attainment and current occupation were collected from guardian by an interviewer and classified according to an updated measure of the Hollingshead index of social status (Barratt, 2012; Hollingshead, 1975). The Hollingshead places education into one of seven categories (1 = “less than 7th grade” to 7 = “graduate/professional training”). Similarly, occupation was measured by nine categories ranked in order of relative prestige (“1= Day laborer, janitor, house cleaner, etc.” to “9= Physician, attorney, professor, etc.”). For all analyses, occupation and education scores were averaged across parents. If neither parent currently worked, they were given a score of zero.

Income-to-needs ratio (INR) was calculated for each participant using the median for a parent-reported household income range divided by the federal poverty threshold for the participant’s household size in that year (US Census Bureau, n.d.). INR is commonly used in SES research and more meaningfully measures the financial resources of a family than just income, such that an INR lower than one represents a family living below the federal poverty threshold.

A public assistance variable was derived as a count of parent-reported enrollment in 10 potential public assistance programs, such as Supplemental Security Income (SSI), Social Security Disability (SSD) and Supplemental Nutrition Assistance Program (SNAP). In follow up analyses, modified forms of the public assistance measure were calculated which measured enrollment exclusively in programs with a low-income requirement (e.g., SSI; i.e., income-restricted public assistance), and conversely for programs without an income-based requirement (e.g., SSD; see Supplement).

To further examine how composite SES scoring might obscure the distinct associations between each SES component and RBA, a composite measure for SES was calculated by averaging the z-score across each SES measure, with public assistance reverse coded since public enrollment has been used elsewhere as a categorical measure of low SES (Kachmar et al., 2019).

### Parent Psychiatric Diagnosis

Guardians self-reported lifetime diagnoses on an extensive list of psychiatric disorders. To control for the potential genetic and environmental influences of parent psychopathology as well as its potential contributions to a reporter bias, we used a measure which captured the number of parents in the home with a psychiatric diagnosis, ranging from zero to two.

### Child Psychiatric Symptoms

The Child Behavior Checklist (CBCL) was administered to parents as a dimensional measure of emotional and behavioral problems (Achenbach, 1994). Our analyses examined age-/sex-normed t-scores for the following subscales: aggressive behavior, anxious/depressed, attention problems, somatic complaints, withdrawn/depressed, rule breaking behavior (oppositional defiant in preschool version). Sensitivity analyses examined whether CBCL associations were specific to the pre-school versus school-age versions of the questionnaire.

### Statistical Analyses

All analyses were performed in R version 4.0.4. Distributions and a correlation matrix for all SES variables are presented in the supplement (Figures S2 – S6). All continuous variables were z-scored per analysis, and we report standardized regression coefficient estimates. Significance tests were two-tailed. Alpha was set at p<.05.

SES components are often intercorrelated. Variance inflation factor threshold (VIF) was employed to assess collinearity amongst independent variables and a threshold was set at 2.5 as a conservative approach to common recommendations (Johnston et al., 2018). If the VIF for a measure exceeded the threshold, collinear variables were removed, and examined with a subset of the independent variables to better understand whether there were confounding effects.

To address our first research question, SES components (occupational prestige, educational attainment, INR, and public assistance) were entered into a multiple linear regression model to examine which components associated with RBA. For comparison, an SES composite was tested in this model in place of its individual components. SES components were also tested individually without the other SES covariates. In follow-up analyses four models examined if chronological age moderated associations between each SES component and RBA. Models covaried for sex assigned at birth, parent psychiatric diagnoses, and scan location.

To address our second research question, we examined associations between RBA and CBCL subscales in linear regression models, controlling for sex and scan location. For symmetry, we also tested parent psychiatric diagnoses as a control in this model. Last, we regressed those CBCL subscales that were significantly associated with RBA on SES components (i.e., occupational prestige, educational attainment, INR, and public assistance), controlling for sex assigned at birth, parent psychiatric diagnoses, and scan location. As SES indicators varied across racial and ethnic groups in this sample, we conducted sensitivity analyses to test both race (i.e., White, African American, Hispanic, Asian, or other; see Supplement) and ethnicity (i.e., Hispanic or not Hispanic) as covariates in all models that include SES components. This served to demonstrate that any differences in RBA or psychiatric symptoms across ethnoracial groups were largely accounted for by inequities in SES.

## RESULTS

### Participant demographics

Demographic information for youth included in this study (N=470) are presented in Table 1. By and large, participants racially identified as White (55%), African American (15%), Hispanic (9%) or belonging to two or more races (17%). Ethnically, 25% of our sample identified as Hispanic. Eighty-six percent of this sample had one or more clinical diagnoses, with neurodevelopmental (63%), anxiety (12.1%) and depressive (5.32%) disorders in highest proportion (Table 1). The median CBCL Total Score was 58.5 (IQR: 50-67; Table 1).

**Table 1.**

|  | **Study Sample**<br>**(N=470)** |
| --- | --- |
| **Birth Sex** |  |
| Male | 288 (61.3%) |
| Female | 182 (38.7%) |
| **Age** |  |
| Mean (SD) | 10.4 (3.26) |
| Median [Min, Max] | 9.88 [5.04, 17.7] |
| **RBA** |  |
| Mean (SD) | -0.0947 (1.57) |
| Median [Min, Max] | -0.194 [-5.88, 4.99] |
| **Hispanic** |  |
| No | 354 (75.3%) |
| Yes | 116 (24.7%) |
| **Race** |  |
| White | 256 (54.5%) |
| African American | 69 (14.7%) |
| Hispanic | 44 (9.4%) |
| Asian | 12 (2.6%) |
| Indian | 1 (0.2%) |
| Native American | 1 (0.2%) |
| Two or more races | 81 (17.2%) |
| Other race | 4 (0.9%) |
| Unknown | 2 (0.4%) |
| **Income-to-Needs Ratio** |  |
| Mean (SD) | 3.95 (2.33) |
| Median [Min, Max] | 3.87 [0.0987, 11.0] |
| **CBCL Internalizing T Score** |  |
| Mean (SD) | 56.9 (11.6) |
| Median [Min, Max] | 58.0 [29.0, 86.0] |
| **CBCL Externalizing T Score** |  |

**Table 1.**
|  | **Study Sample**<br>**(N=470)** |
| --- | --- |
| Mean (SD) | 54.9 (11.9) |
| Median [Min, Max] | 54.0 [29.0, 81.0] |
| **Clinical Diagnosis\*** |  |
| Anxiety Disorders | 57 (12.1%) |
| Depressive Disorders | 25 (5.3%) |
| Disruptive, Impulse Control<br>and Conduct Disorders | 13 (2.8%) |
| Elimination Disorders | 1 (0.2%) |
| Incomplete Evaluation | 5 (1.1%) |
| Neurodevelopmental<br>Disorders | 296 (63.0%) |
| No Diagnosis | 59 (12.6%) |
| Obsessive Compulsive and<br>Related Disorders | 5 (1.1%) |
| Schizophrenia Spectrum and<br>other Psychotic Disorders | 1 (0.2%) |
| Trauma and Stressor Related<br>Disorders | 8 (1.7%) |
| **Parent Psychiatric Diagnoses** |  |
| Neither Parent | 275 (58.5%) |
| One Parent | 157 (33.4%) |
| Both Parents | 38 (8.1%) |

\*Note: Clinical diagnosis consensus was determined by a licensed clinician based on individualized administration of the KSADS-COMP, a semi-structured DSM-5-based psychiatric interview.
SD = Standard Deviation
Min= Minimum
Max = Maximum

On average, families had an income 395% that of the federal poverty threshold, adjusted for household size but not adjusted geographically, and 11% of participants were living below the federal poverty threshold. For reference between 2015 and 2018, New York City’s local poverty measure was between 132-138% that the federal poverty threshold (Unit of the Mayor’s Office for Economic Opportunity, 2018). Roughly half of the parents in this sample did not have a psychiatric diagnosis (59%).

All SES components were moderately correlated (significant Pearson correlations ranged from -0.23 – 0.56 ; Figure S6) but collinearity was sufficiently low to conduct the multiple linear regression analyses (all VIFs < 1.9).

### RBA, Public Assistance, and Parent Occupation

Participants who used less public assistance programs (ß = 0.12, 95%CI: 0.01, 0.23, t_461_ = 2.14, p = .03) and who had lower parental occupation prestige (ß = 0.16, 95%CI: 0.04, 0.28, t_461_ = 2.55, p = .01), also had a lower RBA (Figure 1A-B, Table S1). Neither INR nor parent education significantly associated with RBA (p > 0.5). Notably, participants who reported greater parental psychiatric diagnoses also had a lower RBA (ß = -0.19, 95%CI: -0.33, -0.04, t_461_ = -2.54, p = .01; Figure 1C); the association remained significant when CBCL Total Score was added as an additional control, (ß = -0.19, 95%CI: -0.33, -0.04, t_461_ = -2.49, p = .01). For comparison, SES components were not significant when tested independently without other SES covariates in the model (p > 0.1; Tables S2-S5) and a SES composite was not significant either (p = 0.86; Table S6). Because occupation and public assistance both positively associated with RBA, we retested the SES composite without reverse-coding public assistance, and it was still non-significant (p = 0.12; Table S7). Chronological age did not moderate associations between RBA and any of the SES components (p > .6; Tables S8-S11). Including race and ethnicity in these models had no effect on the results nor were they significant predictors of RBA (Table S12).

**Figure 1.**
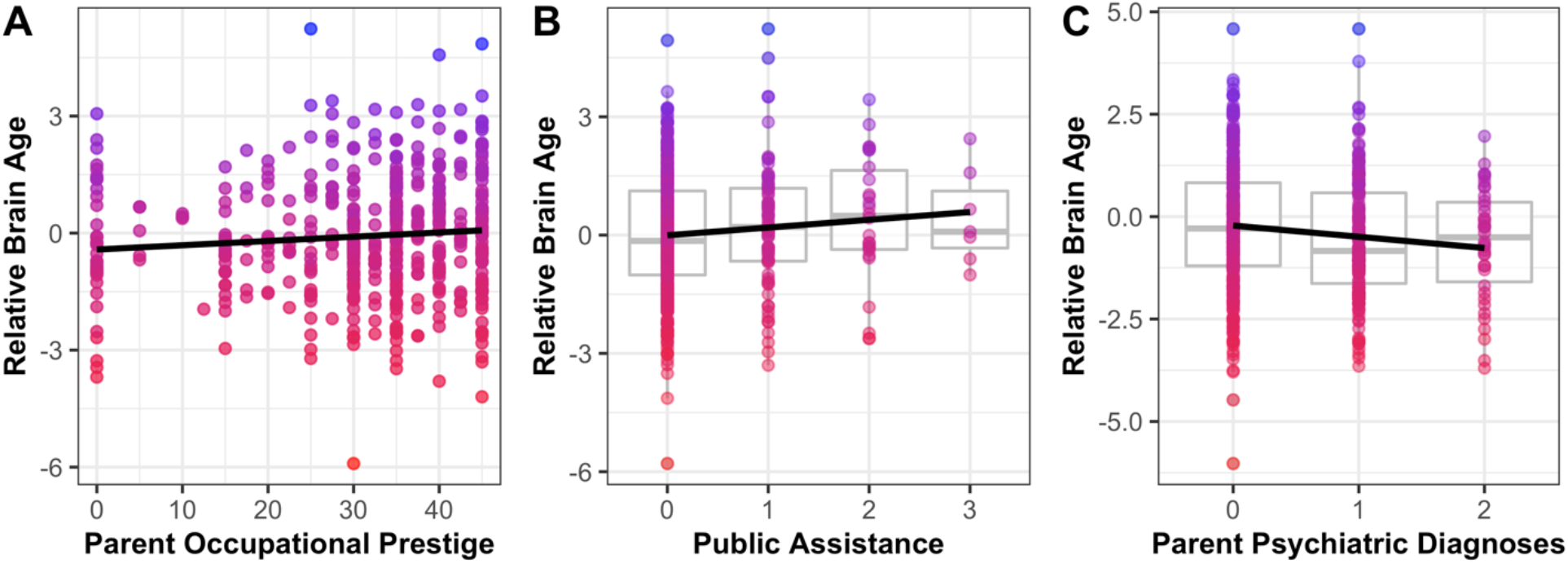
Specific components of socioeconomic status distinctly associated with relative brain age (RBA) when controlling for parent psychiatric diagnosis, sex, and scan location. RBA was positively associated with (A) parent occupational prestige and (B) public assistance, and negatively associated with (C) parent psychiatric diagnoses. RBA was residualized for SES covariates and when plotting the above figures.

To better understand the effects of public assistance on RBA, we parsed this enrollment variable into income-restricted programs and non-income-restricted programs. Participants who specifically used less non-income-restricted public assistance programs had lower RBA (ß = 0.36, 95%CI: 0.10, 0.63, t_461_ = 2.66, p = .01; Table S13). Conversely, enrollment in income-restricted public assistance programs did not associate with RBA (p = .61; Table S14). In a sensitivity analysis, public assistance as a dichotomous variable (any benefit received versus none) was still associated with RBA (ß = 0.27, 95%CI: 0.01, 0.53, t_461_ = 2.00, p = .046; Table S15).

### RBA and CBCL Subscales

Lower RBA significantly associated with greater CBCL anxious/depressed subscale t-scores (ß = -0.09, 95%CI: -0.19, -0.004, t_465_ = -2.05, p = .04), controlling for sex and scan location. No associations were found for other subscales (Tables S16-S21). When parent psychiatric diagnosis was added as a control, the association between RBA and anxious/depressed subscale t-scores was trend level (p = 0.07; Table S22). The RBA x CBCL form type (i.e., school age or preschool version of CBCL) interaction term was not significant (p = .32; Table S23).

### SES Indicators and CBCL Subscales

Participants who had lower parental occupation prestige (ß = -0.15, 95%CI: -0.27, -0.03, t_461_ = -2.38, p = .02), higher parent educational attainment (ß = 0.11, 95%CI: 0.001, 0.216, t_461_ = 1.98, p = .05) and greater parent psychopathology (ß = 0.18, 95%CI: 0.03, 0.32, t_461_ = 2.40, p =.02) also had significantly higher CBCL anxious/depressed subscale scores (Table S24). INR associated with the anxiety/depression subscale at trend-level (ß = -0.10, t_461_ = -1.64, p = .10). Again, in sensitivity analyses, race and ethnicity had no effect on these results (Table S25).

## DISCUSSION

Prior studies have linked specific measures of household SES to differences in youth behavior (Lahelma et al., 2006; Lorant et al., 2003; McLaughlin et al., 2011) and brain structure (Lawson et al., 2013; Noble et al., 2015, 2012; Rakesh et al., 2022), but little is known about household-level SES-related effects on brain predicted aging. Herein, we show that our previously derived SES and psychopathology-blind brain age metrics (Zhao et al., 2019) are in fact associated with specific components of household SES and anxiety/depression symptoms. Low parent occupational prestige and low public assistance program enrollment were both associated with delayed brain aging, which in turn associated with child anxiety/depression symptoms. Further, and consistent with many prior findings (Gilman et al., 2003; Lorant et al., 2003; Pettersson et al., 2019; Power et al., 2007), parent occupational prestige and psychiatric diagnosis were associated with child anxiety/depression symptoms.

### Parental Occupational Prestige

We found that lower parent occupational prestige increases risk for delayed brain aging in children and adolescents. This finding is consistent with prior evidence linking lower SES to alterations in brain structure (Colich et al., 2020; Hanson et al., 2011; Jednoróg et al., 2012; Luby et al., 2013; Mackey et al., 2015; McDermott et al., 2019; Noble et al., 2015, 2012; Ramphal et al., 2021; Ursache et al., 2016) and delays in brain development (Barch et al., 2020; Hair et al., 2015; Hanson et al., 2013). Additionally our finding points to the specificity of parent occupational prestige, above and beyond other household socioeconomic factors on brain age processes, consistent with prior findings showing unique neural correlates of different SES measures (Lawson et al., 2013; Noble et al., 2015, 2012; Rakesh, Zalesky, et al., 2021).

The biological mechanisms and behavioral pathways through which occupational prestige might impact brain aging are not well understood. Parent occupational prestige has been shown to predict toddler cortisol levels (Tarullo et al., 2020), presenting a plausible mechanism for neuroanatomical change via the HPA axis (Burghy et al., 2012; Ellis & Del Giudice, 2019; Merz et al., 2019). Some suggest that occupational prestige shapes parents’ self-worth and motivation (MacKinnon & Langford, 1994), family value systems (Kohn, 1963, 1976; Luster et al., 1989), disciplinary strategies (Parcel & Menaghan, 1994), time spent with child (Roeters et al., 2010), and sources of authority in the home (Kohn, 1959). Brain age may represent a mechanism by which these more proximal home environment factors are biologically embedded in children and adolescents. Future work should aim to understand this mechanism. Importantly, mediators in this process such as parent value systems and disciplinary strategies might be more easily modified than occupation and represent potential targets for intervention.

In our study lower occupational prestige was additionally shown to relate to higher risk for anxiety and depression symptoms. Prior research has shown that occupational prestige associates with concurrent (Kalff et al., 2001) and future (Lombardi & Coley, 2013) behavioral outcomes in youth. More broadly, SES has been shown to predict anxiety (Lähdepuro et al., 2019; McLaughlin et al., 2010) and depression across the lifespan (Gilman et al., 2003; Lorant et al., 2003; Power et al., 2007). Some studies have specifically shown that brain structure (Barch et al., 2020; Noble et al., 2015) and function (Ramphal, Whalen, et al., 2020) mediate the relationship between SES and psychopathology, but none have specifically examined age related processes across the brain. Due to the cross-sectional nature the data, we did not formally test RBA as a mediator in this study. Future work should examine brain age prediction in longitudinal data sets to specifically explore RBA-mediated associations between parent occupational prestige and child depression/anxiety.

### Public Assistance

Public assistance positively associated with brain aging. Surprisingly, enrollment in fewer public assistance programs was associated with more delayed brain aging. In prior studies, public health insurance (Kachmar et al., 2019; Marcin et al., 2003; Ramphal, Whalen, et al., 2020), Supplemental Nutrition Assistance Program, and other forms of public assistance (Kachmar et al., 2019) have been used as proxy measures of low SES and have been linked with adverse outcomes. In contrast, INR only weakly, negatively correlated with public assistance in our study. Indeed, follow-up analyses indicate that the association between public assistance and brain aging was largely driven by programs that did not require low income. Thus, it is unlikely that the effects of low-income are driving our findings. The association between public assistance enrollment and less delayed brain aging likely reflects the benefits associated with public assistance programs despite families having their most basic needs met. It may additionally reflect self-advocacy and the ability to navigate the social safety net system and secure additional resources, a skill that likely benefits a family in more than just this domain. Future work may better explore this construct by including mediators such as parent self-advocacy and testing the above models in a more socioeconomically disadvantaged sample, where basic needs are not met. A more robust understanding of the impact of public assistance programs on child emotional and brain development would allow a more complete appraisal of their efficacy and accessibility.

### Parent Psychiatric Diagnosis

We found a robust association between parent psychopathology and delayed brain aging. In another study, longitudinal decreases in brain aging were observed in youth at familial risk for mood disorders, regardless of the child’s diagnosis (de Nooij et al., 2020). Future studies should aim to disentangle gene-environment interactions whereby parent psychopathology may contribute to child brain development (Burt, 2009; Lahey et al., 2011; Wermter et al., 2010).

Future brain aging research may specifically examine such effects by including more precise measures of psychiatric heritability (e.g., polygenic risk scores) and potential environmental moderators (e.g., parenting practices, dyad relations, socioeconomic markers). Further, epidemiological methods should be employed to account for the potential source of bias in single family homes where psychiatric data is not available for one parent, a limitation in this study.

### Directionality of Aging and Considerations in Brain Age Research

To date, findings on the directionality of risk-associated-aging in youth are equivocal. In one study, an association between an environmental risk composite score and accelerated brain aging was detected, but was not driven meaningfully by parental education and income (Drobinin et al., 2021). Another study found that neighborhood disadvantage was associated with accelerated aging in early adolescence and delayed aging in later adolescence (Rakesh, Cropley, et al., 2021). Across the broader age range included in our study, the association between household SES indicators and brain aging did not vary with age. Discrepancies between study outcomes might derive from differences in how disadvantage was measured. For instance, dissociable neural correlates for neighborhood and household SES have been identified in other work (Gianaros et al., 2017; Miller et al., 2018; Ramphal, DeSerisy, et al., 2020). Recent work with brain aging specifically demonstrated opposing maturational effects for abuse and neglect amongst girls aged 8 to 18 (Keding et al., 2021), a distinction which has was previously explored in a metanalysis comparing neural correlates of threat versus deprivation (McLaughlin et al., 2019).

The conflicting aging directionality highlighted by these studies may also reflect idiosyncrasies across brain age algorithms. Differences in the construction of the age prediction model (e.g., choice of modalities, model, training sample, etc.) can confound interstudy comparison (Lee et al., 2021). These methodological choices must be considered when comparing results across studies, and some researchers emphasize the importance of multidimensional approaches to capture the potentially divergent developmental trajectories of various brain structures (Niu et al., 2021; Rokicki et al., 2021). Alternatively, brain age prediction algorithms may be tailored to the research question (Keding et al., 2021). Still, unitary measures of brain aging maintain the benefit of synthesizing complex information across the brain and may prove useful as a broadly applicable biomarker as these methodological constraints are better understood and addressed.

### Diversity and Brain Age Research

Using a previously derived SES-blind machine learning-based measure of brain aging we investigated the influence of SES on RBA. Because we used this previously validated training set to study this question, our sample size was limited. Recent findings show that cross-validated participant level predictions in neuroimaging research vary with sociodemographic diversity in the training sample (Benkarim et al., 2022). Consistent with these findings we show that parent occupation and public assistance predicted RBA, further suggesting that socioeconomic diversity should be considered in future implementations of brain age prediction algorithms.

## Conclusions

The present study re-examined a previously validated brain age measure in a sample of youth from the HBN dataset through an SES lens. We provide evidence linking SES to delayed brain aging and highlight the importance of measuring the distinct neural and psychopathological correlates of SES components. RBA captured a delayed brain aging phenotype across the brain, which was shared by both low occupational prestige, low public assistance, high familial risk for psychiatric disorders and high anxiety/depression risk for individuals. Such findings point to effects of SES on youth mental health outcomes operating through age-related processes in the brain. Further work with longitudinal data sets will be necessary to better understand the role that aging plays in the development of psychopathology in children and adolescents.

