## Supplemental Materials for "Relative Brain Age Is Associated with Socioeconomic Status and Anxiety/Depression Problems in Youth"

**Supplemental Methods**

**Income-to-Needs Ratio:** The Healthy Brain Network Financial Support Questionnaire interrogated income using 12 bins for household income range. These bins ranged from “Less than $10,000” to “$150,000 or more”, on $10,000 intervals until the penultimate bin which covered a $50,000 range (i.e., “$100,000 to $149,999). Median values for each bin were used in INR calculations, with the exception of the final bin, where $175,000 was chosen as the median value for this group. Income values were then divided by the U.S. Census Bureau’s official weighted poverty threshold, for the participant’s family size in the year the form was administered. The official poverty threshold is not adjusted geographically. The New York City (NYC) poverty measure is between 132-138% that of the federal threshold for 2015-2018, the dates in which income data was collected for this sample. U.S. Census Bureau’s official poverty threshold is more appropriate than the NYC measure for our analyses for a few reasons. U.S. Census Bureau provides a weighted poverty threshold that can be applied to data where the size of the family unit is known but the number of children is not. The NYC measure includes value of near-cash, in-kind benefits such as SNAP, which we are attempting to isolate with our public assistance variable. Lastly, the federal poverty threshold is commonly used in research and could facilitate interstudy comparison.

**Public Assistance:** Public assistance was calculated as sum enrollment in 10 public assistance programs - Supplemental Security Income (SSI), Social Security Disability (SSD), Social Security Retirement Benefits, Social Security Survivors Benefits, Civil Service Pension, Military Retired Pay/Survivors Benefit Plan Annuity (SBP), Medical disability payment, Special Supplemental Nutrition Program for Women, Infants, and Children (WIC), Supplemental Nutrition Assistance Program (SNAP) Benefits, and NYC Cash Assistance Program. Enrollment in just SSI, WIC, SNAP, and the NYC Cash Assistance Program were summed to form the low-income-eligibility public assistance measure. Enrollment in SSD, Social Security Retirement Benefits, Social Security Survivors Benefits, Civil Service Pension, SBP, and Medical disability payment were summed to form the non-low-income eligibility public assistance measure.

**Race**: As Table 1 demonstrates, there were not enough individuals in certain race categories to maintain each level in our analyses. Thus, racial categories “Indian”, “Native American”, “Alaskan Native”, “Pacific Islander”, “Two or more races”, “Other race” and “unknown” were collapsed to one category titled “other” for all analyses.

**Supplemental Tables**

| **Table S1.** Multiple linear regression results for the effects of components of socioeconomic status on relative brain age (RBA) in n = 470 participants, controlling for sex, scan location and parent psychiatric diagnoses. | | | | | |
| --- | --- | --- | --- | --- | --- |
| *Predictors* | *Beta* | *SE* | *95% CI* | *t* | *p* |
| Educational Attainment | -0.01 | 0.06 | -0.12 – 0.10 | -0.19 | 0.850 |
| Occupational Prestige | 0.16 | 0.06 | 0.04 – 0.28 | 2.55 | 0.011 |
| Income-to-Needs Ratio | -0.02 | 0.03 | -0.07 – 0.03 | -0.65 | 0.515 |
| Public Assistance | 0.12 | 0.06 | 0.01 – 0.23 | 2.14 | 0.033 |
| Sex | 0.01 | 0.09 | -0.18 – 0.19 | 0.06 | 0.952 |
| Parent Psychiatric Diagnoses | -0.19 | 0.07 | -0.33 – -0.04 | -2.54 | 0.012 |
| Scan Location [Rutgers] | -0.22 | 0.17 | -0.55 – 0.11 | -1.32 | 0.188 |
| Scan Location [Citigroup] | -0.12 | 0.18 | -0.48 – 0.24 | -0.66 | 0.510 |

| **Table S2.** Multiple linear regression results for the effects of parent educational attainment on relative brain age (RBA) in n = 470 participants, controlling for sex, scan location and parent psychiatric diagnoses. | | | | | |
| --- | --- | --- | --- | --- | --- |
| *Predictors* | *Beta* | *SE* | *95% CI* | *t* | *p* |
| Educational Attainment | 0.03 | 0.05 | -0.07 – 0.12 | 0.58 | 0.564 |
| Sex | -0.01 | 0.10 | -0.20 – 0.18 | -0.10 | 0.920 |
| Parent Psychiatric Diagnoses | -0.16 | 0.07 | -0.30 – -0.02 | -2.21 | 0.028 |
| Scan Location [Rutgers] | -0.23 | 0.17 | -0.56 – 0.09 | -1.40 | 0.161 |
| Scan Location [Citigroup] | -0.16 | 0.18 | -0.51 – 0.20 | -0.85 | 0.394 |

| **Table S3.** Multiple linear regression results for the effects of parent occupational prestige on relative brain age (RBA) in n = 470 participants, controlling for sex, scan location and parent psychiatric diagnoses. | | | | | |
| --- | --- | --- | --- | --- | --- |
| *Predictors* | *Beta* | *SE* | *95% CI* | *t* | *p* |
| Occupational Prestige | 0.08 | 0.05 | -0.02 – 0.17 | 1.64 | 0.102 |
| Sex | -0.01 | 0.09 | -0.19 – 0.18 | -0.07 | 0.948 |
| Parent Psychiatric Diagnoses | -0.16 | 0.07 | -0.30 – -0.02 | -2.21 | 0.028 |
| Scan Location [Rutgers] | -0.26 | 0.17 | -0.59 – 0.06 | -1.58 | 0.114 |
| Scan Location [Citigroup] | -0.18 | 0.18 | -0.53 – 0.17 | -1.00 | 0.319 |

| **Table S4.** Multiple linear regression results for the effects of income-to-needs ratio on relative brain age (RBA) in n = 470 participants, controlling for sex, scan location and parent psychiatric diagnoses. | | | | | |
| --- | --- | --- | --- | --- | --- |
| *Predictors* | *Beta* | *SE* | *95% CI* | *t* | *p* |
| Income-to-Needs Ratio | -0.01 | 0.05 | -0.11 – 0.08 | -0.28 | 0.782 |
| Sex | -0.01 | 0.10 | -0.19 – 0.18 | -0.07 | 0.947 |
| Parent Psychiatric Diagnoses | -0.16 | 0.07 | -0.31 – -0.02 | -2.22 | 0.027 |
| Scan Location [Rutgers] | -0.21 | 0.17 | -0.54 – 0.12 | -1.26 | 0.210 |
| Scan Location [Citigroup] | -0.12 | 0.18 | -0.48 – 0.24 | -0.68 | 0.499 |

| **Table S5.** Multiple linear regression results for the effects of public assistance on relative brain age (RBA) in n = 470 participants, controlling for sex, scan location and parent psychiatric diagnoses. | | | | | |
| --- | --- | --- | --- | --- | --- |
| *Predictors* | *Beta* | *SE* | *95% CI* | *t* | *p* |
| Public Assistance | 0.07 | 0.05 | -0.03 – 0.16 | 1.41 | 0.160 |
| Sex | -0.01 | 0.09 | -0.19 – 0.18 | -0.06 | 0.956 |
| Parent Psychiatric Diagnoses | -0.18 | 0.07 | -0.32 – -0.03 | -2.42 | **0.016** |
| Scan Location [Rutgers] | -0.19 | 0.17 | -0.52 – 0.14 | -1.13 | 0.258 |
| Scan Location [Citigroup] | -0.10 | 0.18 | -0.45 – 0.26 | -0.54 | 0.592 |

| **Table S6.** Multiple linear regression results for the effects of an SES composite score on relative brain age (RBA) in n = 470 participants, controlling for sex, scan location and parent psychiatric diagnoses. The composite measure for SES was calculated by averaging the z-score across each SES measure, with public assistance reverse coded. | | | | | |
| --- | --- | --- | --- | --- | --- |
| *Predictors* | *Beta* | *SE* | *95% CI* | *t* | *p* |
| SES Composite | 0.01 | 0.06 | -0.11 – 0.13 | 0.18 | 0.857 |
| Sex | -0.01 | 0.10 | -0.20 – 0.18 | -0.09 | 0.929 |
| Psychiatric Diagnoses | -0.16 | 0.07 | -0.31 – -0.02 | -2.21 | 0.027 |
| Scan Location [Rutgers] | -0.23 | 0.17 | -0.56 – 0.11 | -1.34 | 0.180 |
| Scan Location [Citigroup] | -0.14 | 0.18 | -0.50 – 0.22 | -0.77 | 0.440 |

| **Table S7.** Multiple linear regression results for the effects of an SES composite score on relative brain age (RBA) in n = 470 participants, controlling for sex, scan location and parent psychiatric diagnoses. The composite measure for SES was calculated by averaging the z-score across each SES measure. | | | | | |
| --- | --- | --- | --- | --- | --- |
| *Predictors* | *Estimates* | *std. Error* | *CI* | *Statistic* | *p* |
| SES Composite | 0.14 | 0.09 | -0.04 – 0.31 | 1.55 | 0.121 |
| Sex | -0.01 | 0.09 | -0.20 – 0.17 | -0.13 | 0.899 |
| Psychiatric Diagnoses | -0.17 | 0.07 | -0.31 – -0.03 | -2.32 | **0.021** |
| Scan Location [Rutgers] | -0.26 | 0.17 | -0.59 – 0.06 | -1.58 | 0.114 |
| Scan Location [Citigroup] | -0.19 | 0.18 | -0.55 – 0.17 | -1.04 | 0.297 |

| **Table S8.** Multiple linear regression results for the age-related effects of parent educational attainment on relative brain age (RBA) in n = 470 participants, controlling for sex, scan location, parent psychiatric diagnoses, and other components of socioeconomic status. | | | | | |
| --- | --- | --- | --- | --- | --- |
| *Predictors* | *Beta* | *SE* | *95% CI* | *t* | *p* |
| Educational Attainment | -0.01 | 0.06 | -0.12 – 0.10 | -0.23 | 0.822 |
| Age | -0.09 | 0.05 | -0.18 – 0.00 | -1.88 | 0.061 |
| Occupational Prestige | 0.14 | 0.06 | 0.02 – 0.27 | 2.28 | **0.023** |
| Income-to-Needs Ratio | -0.04 | 0.06 | -0.16 – 0.08 | -0.67 | 0.503 |
| Public Assistance | 0.09 | 0.06 | -0.02 – 0.20 | 1.57 | 0.116 |
| Sex | 0.00 | 0.10 | -0.18 – 0.19 | 0.02 | 0.982 |
| Scan Location [Rutgers] | -0.21 | 0.17 | -0.55 – 0.12 | -1.27 | 0.205 |
| Scan Location [Citigroup] | -0.13 | 0.18 | -0.50 – 0.23 | -0.73 | 0.467 |
| Educational Attainment * Age | 0.00 | 0.04 | -0.08 – 0.09 | 0.07 | 0.948 |

| **Table S9.** Multiple linear regression results for the age-related effects of parent occupational prestige on relative brain age (RBA) in n = 470 participants, controlling for sex, scan location, parent psychiatric diagnoses, and other components of socioeconomic status. | | | | | |
| --- | --- | --- | --- | --- | --- |
| *Predictors* | *Beta* | *SE* | *95% CI* | *t* | *p* |
| Educational Attainment | -0.01 | 0.06 | -0.12 – 0.10 | -0.21 | 0.831 |
| Occupational Prestige | 0.15 | 0.06 | 0.02 – 0.27 | 2.37 | **0.018** |
| Age | -0.09 | 0.05 | -0.18 – 0.00 | -1.93 | 0.054 |
| Income-to-Needs Ratio | -0.04 | 0.06 | -0.16 – 0.08 | -0.73 | 0.467 |
| Public Assistance | 0.09 | 0.06 | -0.02 – 0.20 | 1.64 | 0.101 |
| Sex | 0.01 | 0.10 | -0.18 – 0.20 | 0.10 | 0.919 |
| Scan Location [Rutgers] | -0.23 | 0.17 | -0.56 – 0.10 | -1.35 | 0.177 |
| Scan Location [Citigroup] | -0.14 | 0.18 | -0.50 – 0.22 | -0.76 | 0.450 |
| Occupational Prestige * Age | -0.05 | 0.05 | -0.14 – 0.04 | -1.06 | 0.288 |

| **Table S10.** Multiple linear regression results for the age-related effects of income-to-needs ratio on relative brain age (RBA) in n = 470 participants, controlling for sex, scan location, parent psychiatric diagnoses, and other components of socioeconomic status. | | | | | |
| --- | --- | --- | --- | --- | --- |
| *Predictors* | *Beta* | *SE* | *95% CI* | *t* | *p* |
| Educational Attainment | -0.01 | 0.06 | -0.12 – 0.10 | -0.25 | 0.799 |
| Occupational Prestige | 0.14 | 0.06 | 0.02 – 0.27 | 2.30 | **0.022** |
| Income-to-Needs Ratio | -0.04 | 0.06 | -0.16 – 0.08 | -0.65 | 0.515 |
| Age | -0.09 | 0.05 | -0.18 – 0.00 | -1.87 | 0.062 |
| Public Assistance | 0.09 | 0.06 | -0.02 – 0.20 | 1.64 | 0.103 |
| Sex | 0.00 | 0.10 | -0.18 – 0.19 | 0.03 | 0.973 |
| Scan Location [Rutgers] | -0.21 | 0.17 | -0.55 – 0.12 | -1.27 | 0.205 |
| Scan Location [Citigroup] | -0.13 | 0.18 | -0.49 – 0.23 | -0.71 | 0.479 |
| INCOME-TO-NEEDS RATIO * Age | -0.02 | 0.05 | -0.11 – 0.07 | -0.52 | 0.604 |

| **Table S11.** Multiple linear regression results for the age-related effects of public assistance on relative brain age (RBA) in n = 470 participants, controlling for sex, scan location, parent psychiatric diagnoses, and other components of socioeconomic status. | | | | | |
| --- | --- | --- | --- | --- | --- |
| *Predictors* | *Beta* | *SE* | *95% CI* | *t* | *p* |
| Educational Attainment | -0.01 | 0.06 | -0.12 – 0.10 | -0.17 | 0.867 |
| Occupational Prestige | 0.14 | 0.06 | 0.02 – 0.26 | 2.24 | 0.026 |
| Income-to-Needs Ratio | -0.04 | 0.06 | -0.16 – 0.08 | -0.67 | 0.502 |
| Public Assistance | 0.08 | 0.06 | -0.03 – 0.19 | 1.47 | 0.142 |
| Age | -0.09 | 0.05 | -0.18 – 0.01 | -1.84 | 0.066 |
| Sex | -0.00 | 0.10 | -0.19 – 0.19 | -0.00 | 0.999 |
| Scan Location [Rutgers] | -0.21 | 0.17 | -0.54 – 0.12 | -1.26 | 0.209 |
| Scan Location [Citigroup] | -0.13 | 0.18 | -0.50 – 0.23 | -0.73 | 0.466 |
| Public Assistance * Age | -0.02 | 0.04 | -0.11 – 0.06 | -0.57 | 0.571 |

| **Table S12**. Multiple linear regression results for the effects of components of socioeconomic status on relative Brain Age (RBA) in n = 470 participants, controlling for sex, scan location, parent psychiatric diagnoses, race, and ethnicity. | | | | | |
| --- | --- | --- | --- | --- | --- |
| *Predictors* | *Beta* | *SE* | *95% CI* | *t* | *p* |
| Educational Attainment | -0.01 | 0.06 | -0.12 – 0.10 | -0.13 | 0.896 |
| Occupational Prestige | 0.16 | 0.06 | 0.03 – 0.28 | 2.51 | **0.012** |
| Income-to-Needs Ratio | -0.05 | 0.06 | -0.17 – 0.07 | -0.77 | 0.443 |
| Public Assistance | 0.12 | 0.06 | 0.01 – 0.23 | 2.16 | **0.031** |
| Sex | 0.00 | 0.10 | -0.18 – 0.19 | 0.03 | 0.976 |
| Scan Location [Rutgers] | -0.24 | 0.17 | -0.58 – 0.09 | -1.42 | 0.155 |
| Scan Location [Citigroup] | -0.13 | 0.18 | -0.49 – 0.24 | -0.69 | 0.492 |
| Psychiatric Diagnoses | -0.19 | 0.07 | -0.34 – -0.04 | -2.57 | **0.011** |
| Hispanic [Yes] | 0.10 | 0.14 | -0.18 – 0.38 | 0.69 | 0.491 |
| Race [Black/African American] | -0.02 | 0.15 | -0.31 – 0.27 | -0.14 | 0.887 |
| Race [Hispanic] | -0.19 | 0.21 | -0.61 – 0.22 | -0.92 | 0.358 |
| Race [Asian] | -0.34 | 0.30 | -0.92 – 0.25 | -1.12 | 0.263 |
| Race [Other] | -0.17 | 0.14 | -0.45 – 0.10 | -1.24 | 0.216 |

| **Table S13**. Multiple linear regression results for the effects of components of socioeconomic status on relative Brain Age (RBA) in n = 470 participants, controlling for sex, scan location, and parent psychiatric diagnoses. A modified form of the public assistance measure using just those without an income-based requirement was used in this model. | | | | | |
| --- | --- | --- | --- | --- | --- |
| *Predictors* | *Beta* | *SE* | *95% CI* | *t* | *p* |
| Educational Attainment | -0.01 | 0.06 | -0.12 – 0.09 | -0.26 | 0.798 |
| Occupational Prestige | 0.15 | 0.06 | 0.03 – 0.27 | 2.51 | **0.012** |
| Income-to-Needs Ratio | -0.07 | 0.06 | -0.18 – 0.04 | -1.21 | 0.228 |
| Non-Income-Restricted Public Assistance | 0.36 | 0.14 | 0.10 – 0.63 | 2.66 | **0.008** |
| Sex | 0.00 | 0.09 | -0.18 – 0.19 | 0.02 | 0.986 |
| Scan Location [Rutgers] | -0.25 | 0.17 | -0.58 – 0.08 | -1.49 | 0.137 |
| Scan Location [Citigroup] | -0.15 | 0.18 | -0.51 – 0.21 | -0.83 | 0.410 |
| Psychiatric Diagnoses | -0.17 | 0.07 | -0.31 – -0.03 | -2.34 | **0.020** |

| **Table S14**. Multiple linear regression results for the effects of components of socioeconomic status on relative Brain Age (RBA) in n = 470 participants, controlling for sex, scan location, and parent psychiatric diagnoses. A modified form of the public assistance measure counting just those with an income-based requirement was used in this model. | | | | | |
| --- | --- | --- | --- | --- | --- |
| *Predictors* | *Beta* | *SE* | *95% CI* | *t* | *p* |
| Educational Attainment | 0.00 | 0.06 | -0.11 – 0.11 | 0.01 | 0.991 |
| Occupational Prestige | 0.13 | 0.06 | 0.01 – 0.24 | 2.06 | 0.040 |
| Income-to-Needs Ratio | -0.07 | 0.06 | -0.19 – 0.05 | -1.16 | 0.248 |
| Income-Restricted Public Assistance | 0.05 | 0.10 | -0.15 – 0.25 | 0.51 | 0.611 |
| Sex | 0.01 | 0.10 | -0.18 – 0.19 | 0.05 | 0.957 |
| Scan Location [Rutgers] | -0.23 | 0.17 | -0.56 – 0.10 | -1.36 | 0.176 |
| Scan Location [Citigroup] | -0.13 | 0.18 | -0.50 – 0.23 | -0.73 | 0.465 |
| Psychiatric Diagnoses | -0.16 | 0.07 | -0.31 – -0.02 | -2.22 | 0.027 |

| **Table S15**. Multiple linear regression results for the effects of components of socioeconomic status on relative Brain Age (RBA) in n = 470 participants, controlling for sex, scan location, and parent psychiatric diagnoses. A dichotomized form of the public assistance measure measuring none or any use of public assistance was implemented in this model. | | | | | |
| --- | --- | --- | --- | --- | --- |
| *Predictors* | *Beta* | *SE* | *95% CI* | *t* | *p* |
| Educational Attainment | -0.00 | 0.05 | -0.11 – 0.11 | -0.03 | 0.973 |
| Occupational Prestige | 0.15 | 0.06 | 0.03 – 0.27 | 2.43 | **0.015** |
| Income-to-Needs Ratio | -0.04 | 0.06 | -0.16 – 0.07 | -0.72 | 0.469 |
| Public Assistance [Yes] | 0.27 | 0.13 | 0.01 – 0.53 | 2.00 | **0.046** |
| Sex | 0.01 | 0.09 | -0.18 – 0.19 | 0.09 | 0.931 |
| Psychiatric Diagnoses | -0.18 | 0.07 | -0.33 – -0.04 | -2.46 | **0.014** |
| Scan Location [Rutgers] | -0.22 | 0.17 | -0.55 – 0.11 | -1.31 | 0.191 |
| Scan Location [Citigroup] | -0.12 | 0.18 | -0.48 – 0.24 | -0.64 | 0.521 |

| **Table S16**. Multiple linear regression results for the effects of relative brain age (RBA) on aggressive behavior subscale t-scores in n = 470 participants, controlling for sex and scan location. | | | | | |
| --- | --- | --- | --- | --- | --- |
| *Predictors* | *Beta* | *SE* | *95% CI* | *t* | *p* |
| RBA | -0.02 | 0.05 | -0.11 – 0.07 | -0.43 | 0.666 |
| Sex | -0.01 | 0.10 | -0.20 – 0.17 | -0.15 | 0.881 |
| Scan Location [Rutgers] | -0.11 | 0.17 | -0.44 – 0.21 | -0.69 | 0.489 |
| Scan Location [Citigroup] | -0.12 | 0.18 | -0.47 – 0.24 | -0.66 | 0.512 |

| **Table S17**. Multiple linear regression results for the effects of relative brain age (RBA) on anxiety/depression subscale t-scores in n = 470 participants, controlling for sex and scan location. | | | | | |
| --- | --- | --- | --- | --- | --- |
| *Predictors* | *Beta* | *SE* | *95% CI* | *t* | *p* |
| RBA | -0.09 | 0.05 | -0.19 – -0.00 | -2.05 | 0.041 |
| Sex | 0.04 | 0.09 | -0.14 – 0.23 | 0.44 | 0.662 |
| Scan Location [Rutgers] | -0.12 | 0.16 | -0.44 – 0.21 | -0.71 | 0.481 |
| Scan Location [Citigroup] | -0.15 | 0.18 | -0.50 – 0.20 | -0.85 | 0.395 |

| **Table S18**. Multiple linear regression results for the effects of relative brain age (RBA) on attention problems subscale t-scores in n = 470 participants, controlling for sex and scan location. | | | | | |
| --- | --- | --- | --- | --- | --- |
| *Predictors* | *Beta* | *SE* | *95% CI* | *t* | *p* |
| RBA | 0.01 | 0.05 | -0.08 – 0.10 | 0.15 | 0.878 |
| Sex | 0.15 | 0.10 | -0.04 – 0.34 | 1.56 | 0.118 |
| Scan Location [Rutgers] | 0.07 | 0.17 | -0.25 – 0.40 | 0.43 | 0.666 |
| Scan Location [Citigroup] | 0.10 | 0.18 | -0.25 – 0.45 | 0.57 | 0.572 |

| **Table S19**. Multiple linear regression results for the effects of relative brain age (RBA) on somatic complaints subscale t-scores in n = 470 participants, controlling for sex and scan location. | | | | | |
| --- | --- | --- | --- | --- | --- |
| *Predictors* | *Beta* | *SE* | *95% CI* | *t* | *p* |
| RBA | -0.01 | 0.05 | -0.10 – 0.08 | -0.29 | 0.774 |
| Sex | 0.23 | 0.09 | 0.05 – 0.42 | 2.50 | 0.013 |
| Scan Location [Rutgers] | -0.29 | 0.16 | -0.61 – 0.04 | -1.75 | 0.081 |
| Scan Location [Citigroup] | -0.42 | 0.18 | -0.77 – -0.07 | -2.37 | 0.018 |

| **Table S20**. Multiple linear regression results for the effects of relative brain age (RBA) on withdrawn/depressed subscale t-scores in n = 470 participants, controlling for sex and scan location. | | | | | |
| --- | --- | --- | --- | --- | --- |
| *Predictors* | *Beta* | *SE* | *95% CI* | *t* | *p* |
| RBA | -0.02 | 0.05 | -0.11 – 0.07 | -0.37 | 0.715 |
| Sex | -0.01 | 0.10 | -0.19 – 0.18 | -0.06 | 0.953 |
| Scan Location [Rutgers] | 0.08 | 0.17 | -0.24 – 0.41 | 0.49 | 0.627 |
| Scan Location [Citigroup] | 0.04 | 0.18 | -0.31 – 0.39 | 0.22 | 0.829 |

| **Table S21**. Multiple linear regression results for the effects of relative brain age (RBA) on rule-breaking behavior subscale t-scores in n = 470 participants, controlling for sex and scan location. | | | | | |
| --- | --- | --- | --- | --- | --- |
| *Predictors* | *Beta* | *SE* | *95% CI* | *t* | *p* |
| RBA | 0.05 | 0.05 | -0.04 – 0.15 | 1.17 | 0.243 |
| Sex | 0.14 | 0.09 | -0.05 – 0.32 | 1.43 | 0.154 |
| Scan Location [Rutgers] | -0.04 | 0.17 | -0.36 – 0.29 | -0.23 | 0.820 |
| Scan Location [Citigroup] | -0.02 | 0.18 | -0.37 – 0.33 | -0.11 | 0.914 |

| **Table S22**. Multiple linear regression results for the effects of relative brain age (RBA) on anxiety/depression subscale t-scores in n = 470 participants, controlling for sex, scan location and parent psychiatric diagnoses. | | | | | |
| --- | --- | --- | --- | --- | --- |
| *Predictors* | *Beta* | *SE* | *95% CI* | *t* | *p* |
| RBA | -0.08 | 0.05 | -0.18 – 0.01 | -1.83 | 0.068 |
| Sex | 0.03 | 0.09 | -0.16 – 0.21 | 0.27 | 0.789 |
| Scan Location [Rutgers] | -0.08 | 0.17 | -0.40 – 0.25 | -0.47 | 0.640 |
| Scan Location [Citigroup] | -0.15 | 0.18 | -0.50 – 0.20 | -0.82 | 0.412 |
| Psychiatric Diagnoses | 0.16 | 0.07 | 0.01 – 0.30 | 2.14 | 0.033 |

| **Table S23**. Multiple linear regression results for the effects of relative brain age (RBA) on anxiety/depression subscale t-scores with a moderator for form type (pre-school v. school-age) in n = 470 participants, controlling for sex, and scan location. | | | | | |
| --- | --- | --- | --- | --- | --- |
| *Predictors* | *Beta* | *SE* | *95% CI* | *t* | *p* |
| RBA | -0.09 | 0.05 | -0.18 – 0.01 | -1.82 | 0.069 |
| CBCL form [pre] | -0.25 | 0.19 | -0.63 – 0.12 | -1.32 | 0.189 |
| Sex | 0.04 | 0.09 | -0.14 – 0.23 | 0.44 | 0.662 |
| Scan Location [Rutgers] | -0.10 | 0.17 | -0.42 – 0.23 | -0.58 | 0.561 |
| Scan Location [Citigroup] | -0.14 | 0.18 | -0.49 – 0.22 | -0.76 | 0.446 |
| RBA * CBCL form [pre-school] | -0.25 | 0.25 | -0.75 – 0.25 | -0.99 | 0.320 |

| **Table S24**. Multiple linear regression results for the effects of relative brain age (RBA) on anxiety/depression subscale t-scores in n = 470 participants, controlling for sex and scan location. | | | | | |
| --- | --- | --- | --- | --- | --- |
| *Predictors* | *Beta* | *SE* | *95% CI* | *t* | *p* |
| Educational Attainment | 0.11 | 0.05 | 0.00 – 0.22 | 1.98 | 0.048 |
| Occupational Prestige | -0.15 | 0.06 | -0.27 – -0.03 | -2.38 | 0.018 |
| Income-to-Needs Ratio | -0.10 | 0.06 | -0.22 – 0.02 | -1.64 | 0.102 |
| Public Assistance | -0.01 | 0.06 | -0.12 – 0.10 | -0.12 | 0.904 |
| Sex | 0.03 | 0.09 | -0.16 – 0.21 | 0.29 | 0.771 |
| Psychiatric Diagnoses | 0.18 | 0.07 | 0.03 – 0.32 | 2.40 | 0.017 |
| Scan Location [Rutgers] | 0.03 | 0.17 | -0.29 – 0.36 | 0.20 | 0.841 |
| Scan Location [Citigroup] | -0.06 | 0.18 | -0.41 – 0.30 | -0.30 | 0.761 |

| **Table S25**. Multiple linear regression results for the effects of relative brain age (RBA) on anxiety/depression subscale t-scores in n = 470 participants, controlling for sex, scan location, race and ethnicity. | | | | | |
| --- | --- | --- | --- | --- | --- |
| *Predictors* | *Beta* | *SE* | *95% CI* | *t* | *p* |
| Educational Attainment | 0.09 | 0.06 | -0.02 – 0.20 | 1.62 | 0.106 |
| Occupational Prestige | -0.14 | 0.06 | -0.26 – -0.02 | -2.24 | 0.025 |
| Income-to-Needs Ratio | -0.11 | 0.06 | -0.23 – 0.01 | -1.83 | 0.068 |
| Public Assistance | 0.02 | 0.06 | -0.09 – 0.13 | 0.27 | 0.785 |
| Sex | 0.04 | 0.09 | -0.14 – 0.23 | 0.45 | 0.652 |
| Psychiatric Diagnoses | 0.16 | 0.07 | 0.02 – 0.31 | 2.24 | 0.026 |
| Scan Location [Rutgers] | 0.03 | 0.17 | -0.31 – 0.36 | 0.17 | 0.868 |
| Scan Location [Citigroup] | -0.04 | 0.18 | -0.39 – 0.32 | -0.19 | 0.846 |
| Hispanic [Yes] | -0.08 | 0.14 | -0.36 – 0.20 | -0.58 | 0.565 |
| Race [Black/African American] | -0.17 | 0.14 | -0.45 – 0.11 | -1.19 | 0.234 |
| Race [Hispanic] | -0.18 | 0.21 | -0.60 – 0.23 | -0.88 | 0.379 |
| Race [Asian] | 0.54 | 0.30 | -0.04 – 1.12 | 1.82 | 0.069 |
| Race [Other] | -0.12 | 0.14 | -0.39 – 0.16 | -0.84 | 0.403 |

**Supplemental Figures**

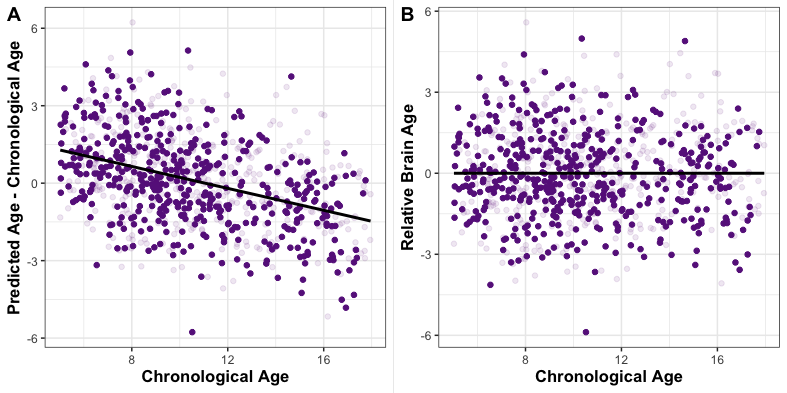

Figure S1. Relative Brain Age as a Corrective Measure. (A) The brain age gap (Predicted – Chronological Age) overestimates aging in younger participants, and underestimates aging in older participants. (B) Regressing predicted age on chronological age and saving the RBAuals eliminates this bias.

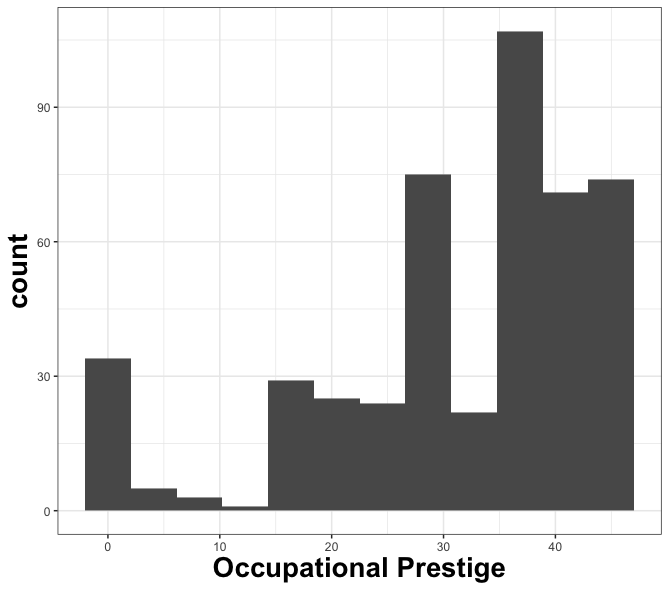

Figure S3. Distribution for parent occupational prestige as measured by the Hollingshead Index

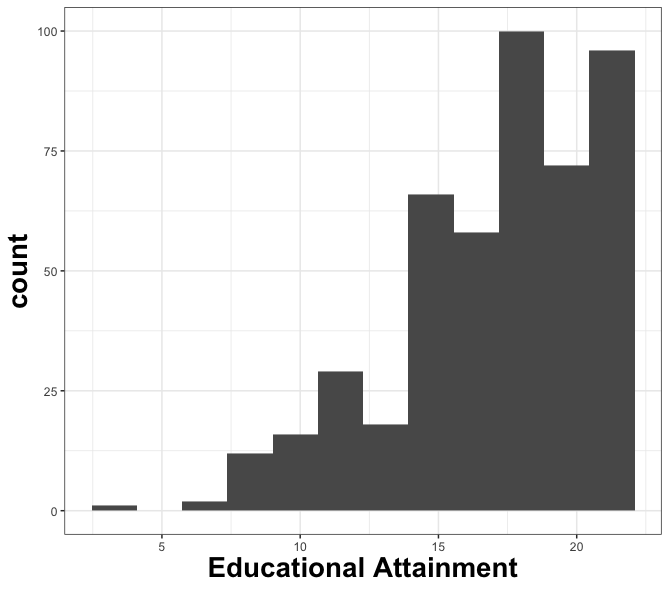

Figure S2. Distribution for parent educational attainment as measured by the Hollingshead Index

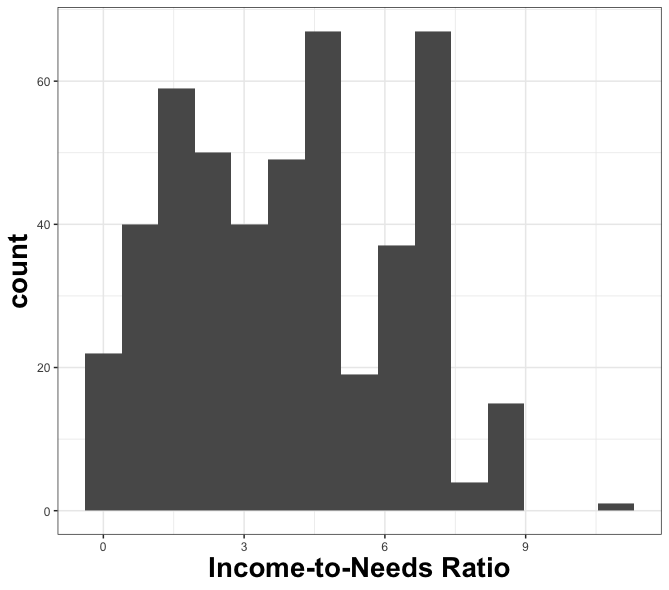

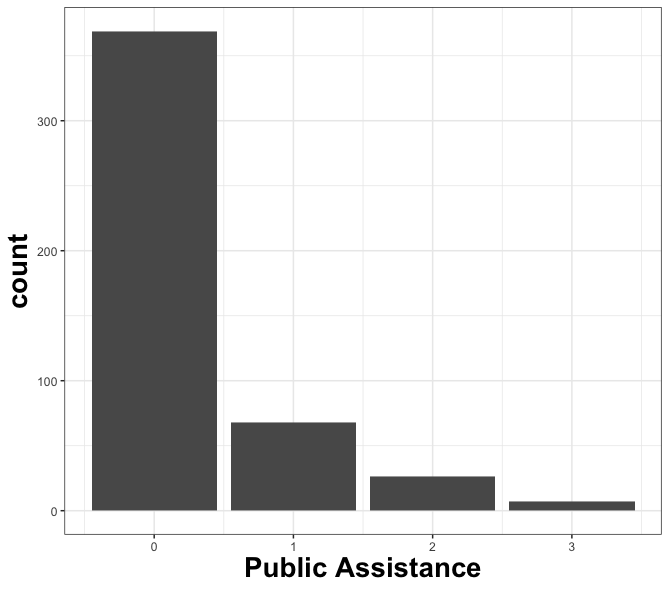

Figure S5. Distribution for enrollment in public assistance programs

Figure S4. Distribution for income-to-needs ratio, calculated using the U.S. Census Bureau official poverty threshold

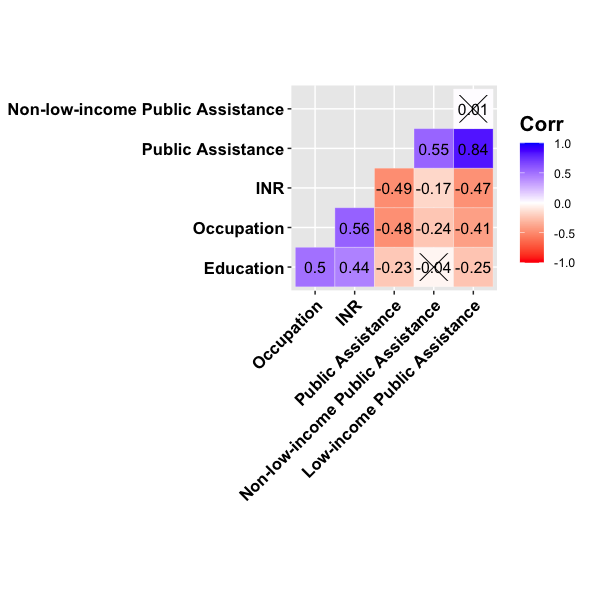

Figure S6. Pearson correlation coefficients for SES component measures. “X” indicates correlation is not significant at α = 0.5.
